# Multi-omics profiling of collagen-induced arthritis mouse model reveals early metabolic dysregulation via SIRT1 axis

**DOI:** 10.1101/2022.03.09.483621

**Authors:** Lingzi Li, Janina Freitag, Christian Asbrand, Bogdan Munteanu, Bei-Tzu Wang, Ekaterina Zezina, Michel Didier, Gilbert Thill, Corinne Rocher, Matthias Herrmann, Nadine Biesemann

## Abstract

Rheumatoid arthritis (RA) is characterized by joint infiltration of immune cells and synovial inflammation which leads to progressive disability. Current treatments improve the disease outcome, but the unmet medical need is still high. New discoveries over the last decade have revealed the major impact of cellular metabolism on immune cell functions. So far, a comprehensive understanding of metabolic changes during disease development, especially in the diseased microenvironment, is still limited. Therefore, we studied the longitudinal metabolic changes during the development of murine arthritis integrating metabolomics and bulk RNA-seq data. We identified an early change in macrophage pathways which was accompanied by oxidative stress, a drop in NAD+ level and induction of glucose transporters. We discovered inhibition of SIRT1, a NAD-dependent histone deacetylase and confirmed its dysregulation in human macrophages and synovial tissue of RA patients. Mining this database should enable the discovery of novel metabolic targets and therapy opportunities in RA.

## Introduction

Rheumatoid arthritis (RA) is a chronic autoimmune disease characterized by infiltration of immune cells in joints, synovial hyperplasia, and joint damage [1]. The etiology of RA is multifactorial, which involves risk factors of genetics and environment [2]. As RA patients exhibit broad biological heterogeneity with varying clinical outcomes [3], the underlying molecular mechanism of RA onset and manifestation remains elusive by far. Deeper insights into the early stages of the disease are urgently needed to develop novel therapies that break the ceiling of efficacy of currently approved drugs.

Immunometabolism has emerged in the recent years as a potential mechanism central to the regulation of immune cells [4]. Major aspects of metabolic reprogramming in pro-inflammatory immune cells include a switch to aerobic glycolysis to support rapid ATP production and nucleic acid synthesis [5]. On the other hand, anti-inflammatory cells tend to rely on oxidative phosphorylation (OXPHOS) and fatty acid oxidation for efficient energy production to sustain longevity [6]. There is growing evidence that metabolism plays a critical role in RA. Joints from RA patients have increased uptake of glucose, along with upregulation of glycolysis in fibroblast-like synoviocytes (FLS) [7–9]. We have recently described induction of glycolytic genes and hypoxia-induced factor 1 alpha (HIF1A) after TNF (tumor necrosis factor alpha) stimulation of RA-FLS [10]. Furthermore, excessive production of reactive oxygen species (ROS) and oxidative stress are implicated in the pathogenesis of arthritis joints [11–13]. However, little is known about the impact of cellular metabolism in the evolution of RA pathogenesis.

Here we present a multi-omics approach to fill the knowledge gap and to provide a comprehensive overview on the longitudinal metabolic changes over the course of RA progression. We utilized the collagen-induced arthritis (CIA) mouse model, which resembles the dynamic pathological features of the acute phase of RA [14, 15]. We combined bulk RNA-seq and metabolomics on CIA mouse paws with metabolomics on CIA plasma, and back-translated our findings with clinical data from different RA disease stages. In addition, the longitudinal plasma metabolomics dataset offers a resource for mining early RA biomarkers. Finally, we identified SIRT1 axis in macrophages as a potential driver of early RA disease stages.

## Results

### Comprehensive characterization of arthritis progression in CIA mice

CIA was induced by two immunizations with Type II collagen in Complete Freund’s adjuvant (CFA) at day zero, and with Type II collagen in Incomplete Freund’s adjuvant (IFA) at day twenty-one. Arthritis development was followed over 10 weeks and mice were sacrificed for tissue collection at eight different time points covering early, middle and late disease stages (Fig. 1a). Evaluation of arthritis was conducted by scoring based on the swelling and redness of the paws (Fig. 1b). Score 0 indicated no visible sign of inflammation. The first sign of paw inflammation (arthritis score > 0) occurred 2 weeks after the 1st immunization and the disease severity progressed over the course of the whole study (Fig. 1c). Variation in arthritis severity was observed (Fig. 1d), matching to previous studies on CIA mouse models [16, 17]. The arthritis manifestation was accompanied by splenomegaly that became evident from week two on (Fig. 1e), which indicates hyperplasia of immune cells. Plasma pro-inflammatory cytokine TNF-α (Fig. 1f), neutrophil-recruiting chemokine CXCL1 (Fig. 1g) and monocyte-recruiting chemokine CCL5 (Fig. 1h) showed pronounced increases already at day two. Elevation of TNF-α level in plasma was consistent over the course of the study, whereas chemokines CXCL1 and CCL5 increased transiently in response to the immunizations. As TNF-α is highly involved in inflammation and osteoclastogenesis in human RA [18], we reasoned that pathogenesis of arthritis already started in CIA mice as early as day two.

**Fig. 1:**
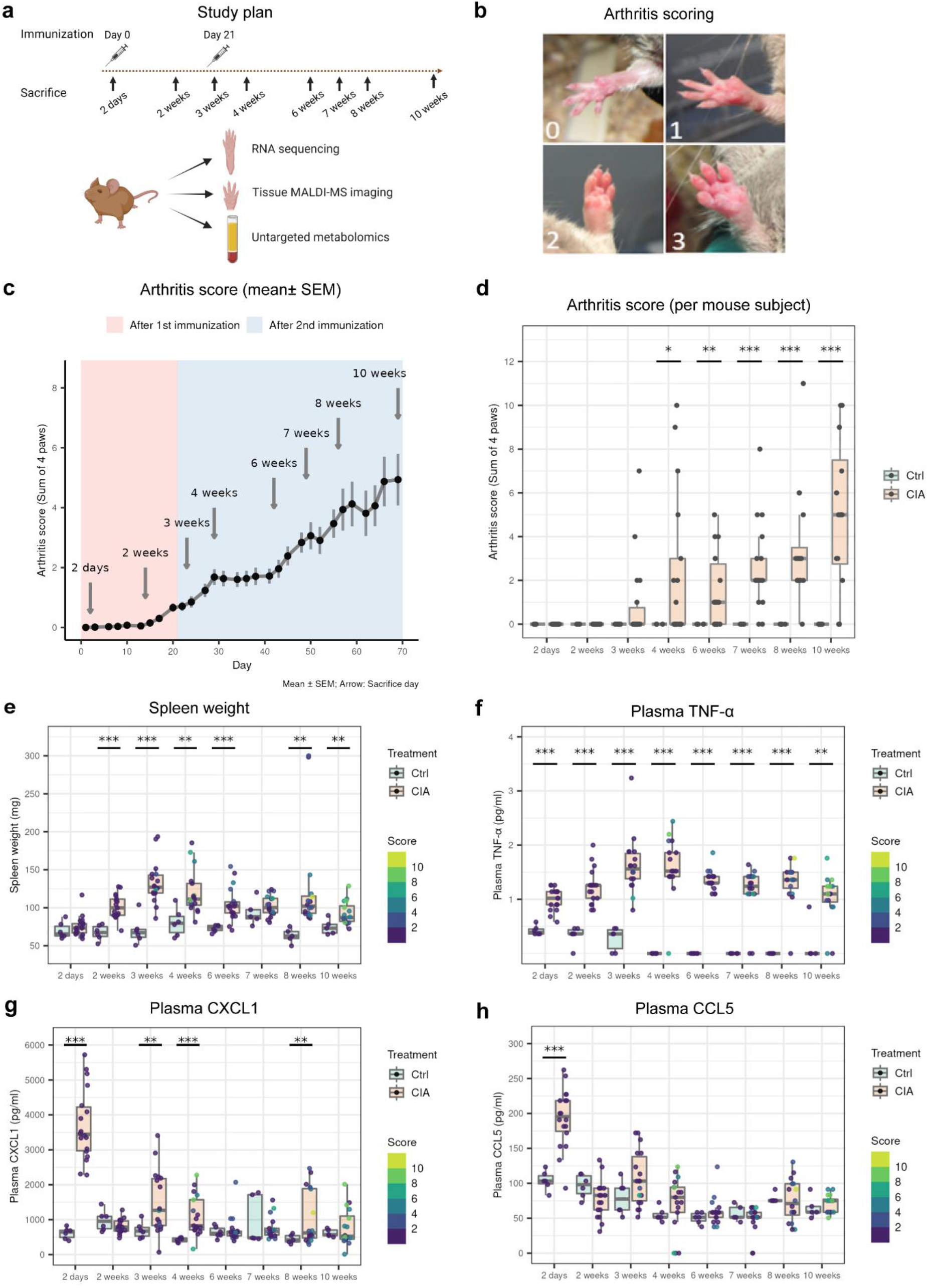
Arthritis progression in CIA mice. **a** Experimental scheme. **b** Representative mouse paws with corresponding arthritis scores. **c** Evolution of arthritis severity by scores (sum of 4 paws, plotted as mean ± SEM). **d** Arthritis severity at each sacrifice time point per mouse subject. **e** Spleen weight measured at each sacrifice time point. **f** Plasma TNF-α concentration. **g** Plasma CXCL1 concentration. **h** Plasma CCL5 concentration. The cytokines and chemokines were measured by mouse magnetic Luminex assays. The dots on the boxplots represent individual mouse samples. At each time point, the sample size for controls is ≥5 and the sample size for CIA is ≥ 16. The color of each dot represents the arthritis score of the mouse (sum of 4 paws). Two-tail Welch t tests were performed (p < 0.05 *, p < 0.01 **, p < 0.001 ***).

### Early metabolic changes in CIA mouse plasma

In addition to the observed changes on cytokines and chemokines, we wanted to investigate how the metabolome changes over the course of arthritis progression. Therefore, CIA plasma samples were analyzed using an untargeted metabolomics approach with ultra high performance liquid chromatography-tandem mass spectroscopy (UPLC-MS/MS). Principal component analysis (PCA) revealed a partial separation of the CIA samples from the control samples starting from week three (Fig. 2a). The UPLC-MS/MS analysis detected 929 metabolites (Fig. 2b) and surprisingly, there were already over 100 significantly altered metabolites at day two (Fig. 2c). A large share of changes in the early time points came from peptides, energy metabolites, cofactors and vitamins (Fig. 2d). Furthermore, peptides and energy metabolites tended to be more upregulated than downregulated, whereas cofactors and vitamins tended to be more downregulated (Fig. 2e).

**Fig. 2:**
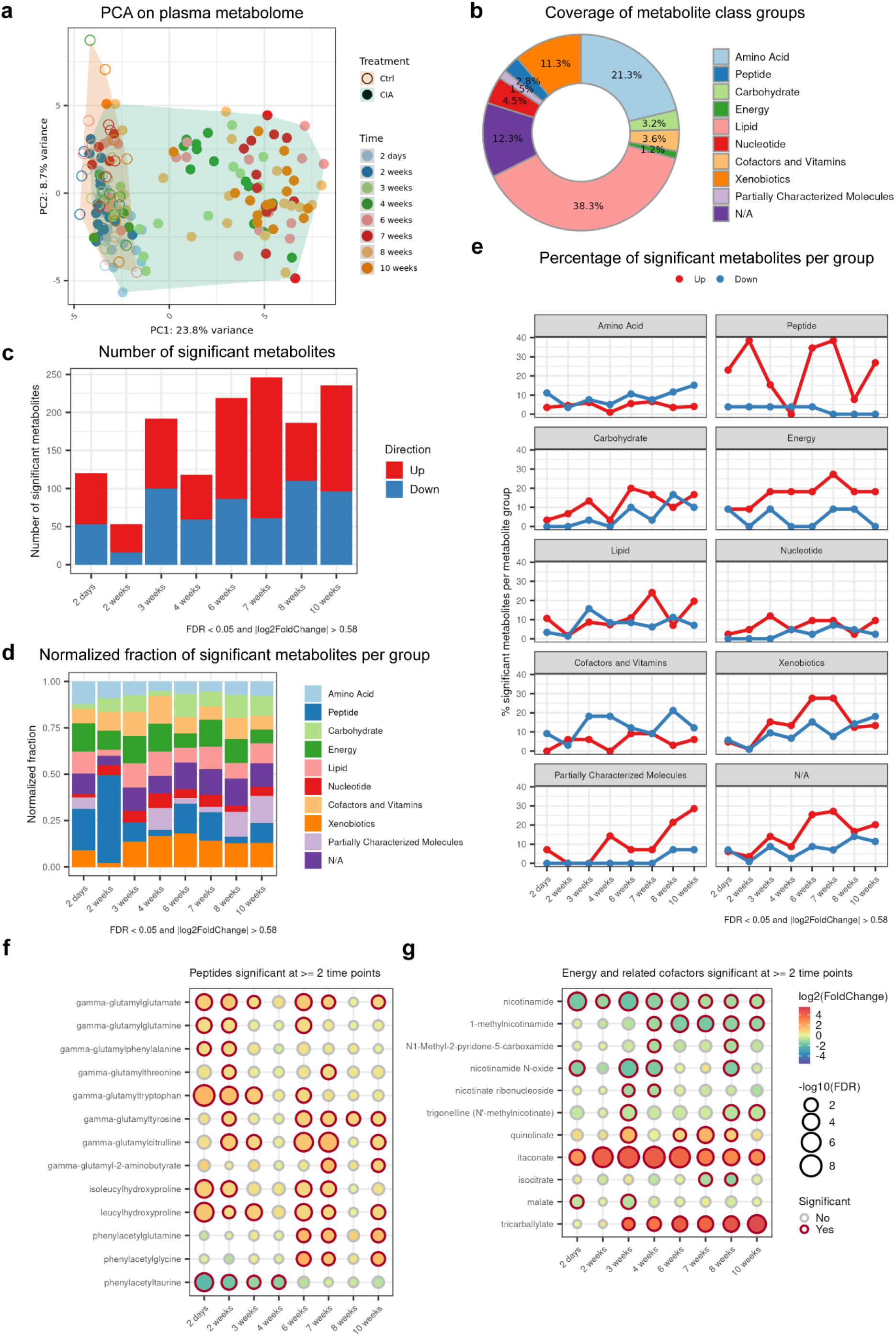
UPLC-MS/MS metabolomics revealed early metabolic alterations in plasma of CIA mice. UPLC-MS/MS metabolomics in total detected 929 metabolites. At each time point, the sample size for controls is ≥5 and the sample size for CIA is ≥ 16. **a** Principal component analysis (PCA) score plot with plasma samples projected onto the first 2 principal components. **b** Percentage of each metabolic group detected by UPLC-MS/MS. **c** Number of significant metabolites at each time point (CIA samples compared to the controls). Significance was decided by cutoffs of FDR < 0.05 and fold change > 1.5 in up or down direction. **d** Percentage of significant metabolites per metabolite group at each time point. Counts of significant metabolites were first normalized to the total number of detected metabolites per metabolite group, and then assembled into a percent stacked bar plot. **e** Percentage of upregulated and downregulated significant metabolites per group over time. **f-g** Fold changes and significances of peptides, energy metabolites and related cofactors that are significant at >= 2 time points (these 2 figures were plotted on the same scale). Two-way ANOVA was done with multiple testing correction using false discovery rate (FDR) method. Significance was decided by cutoffs of FDR < 0.05 and fold change > 1.5 in up or down direction.

We then proceeded to investigate the metabolite molecules that drove these changes (Fig. 2f). Of the peptides that are significantly altered in at least 2 time points, the group of gamma-glutamyl amino acids are significantly increased in early time points (day two and week two, Fig. 2f). These early perturbations persisted until the last time point (week ten) for gamma-glutamylglutamate, gamma-glutamyltyrosine and gamma-glutamylcitrulline (Fig. 2f). The elevated plasma gamma-glutamyl amino acids are a sign of increased activities in the gamma-glutamyl cycle [19], which is essential for metabolism of the antioxidant glutathione and re-utilization of amino acids. Gamma-glutamyl transferase (GGT) on the cell membranes degrades extracellular glutathione into dipeptide cysteinyl-glycine and transfers gamma-glutamyl moiety to amino acids to form gamma-glutamyl amino acids. These are taken up by the cells and re-utilized as precursors for the intracellular synthesis of glutathione and proteins to support cellular functions [20]. Glutathione is the central metabolite in this pathway and has multiple functions including defense against oxidative stress, redox signaling and cell proliferation [21].

The evidence of dysregulated redox signaling was further strengthened by early changes in nicotinamide (Fig. 2g). Among the energy metabolites and the related cofactors, nicotinamide and itaconate had persistent perturbations in all 8 time points from day two until week ten (Fig. 2g). Nicotinamide belongs to the classical NAD+ salvage pathway, where nicotinamide is recycled back into NAD+ [22]. Besides, NAD+ can also be converted from nicotinic acid through the Preiss-Handler pathway [23, 24]. We observed reduction in nicotinamide and nicotinamide N-oxide in the early time point at day two (Fig. 2g). Other nicotinamide metabolites (1-methylnicotinamide and N1-Methyl-2-pyridone-5-carboxamide) and precursors of NAD+ in the Preiss-Handler pathway (nicotinate ribonucleoside and trigonelline) began to show significant reduction at later time points (week three - four). Interestingly, the level of a precursor for the NAD+ de novo biosynthesis, quinolinate, started to rise from week three. Although the current plasma untargeted metabolomics analysis did not profile NAD+, we reasoned that the plasma NAD+ level is probably depleted in CIA mice in early arthritis, resulting in decreased levels of nicotinamide metabolites in its salvage pathway.

The robust surge in itaconate from day two onwards pointed towards an early shift in the tricarboxylic acid (TCA) cycle (Fig. 2g). Itaconate is produced by diverting cis-aconitate away from the TCA cycle upon pro-inflammatory macrophage activation [25]. This also explains the observation that the downstream molecules of TCA cycle, isocitrate and malate, decreased in plasma of CIA mice. In addition, these changes were in parallel to early rises in plasma TNF-α (Fig. 1f) which is mainly secreted from macrophages. This result supports the hypothesis that activation of pro-inflammatory macrophage plays a key role in early pathogenesis of RA.

### Transcriptome analysis identified SIRT1 as upstream regulators

Next, we performed bulk RNA-seq on the paw homogenates to study gene expression over disease progression. Paw transcriptome PCA showed a gradual separation between samples as the arthritis score increased (Fig. 3a). The samples from visually healthy paws (arthritis score = 0) tended to cluster on the lower left part of the PCA plot, whereas inflamed samples tended to move to the upper right part as the arthritis scores increased. When comparing transcriptomes between different time points (Fig. 3b), we saw a partial separation of the CIA samples from the controls starting from week three, matching the pattern on the PCA plot of CIA plasma metabolome (Fig. 2a). Taken together, this further confirmed the phenotype heterogeneity of CIA mice. Compared to the plasma metabolomics with >100 significant metabolites at the early time point at day two, the paw transcriptomics had only a handful of significant genes at this time point. Later, a sharp increase of significant differentially expressed genes occurred at week three. Furthermore, the majority of expression changes were upregulated genes (Fig. 3c).

**Fig. 3:**
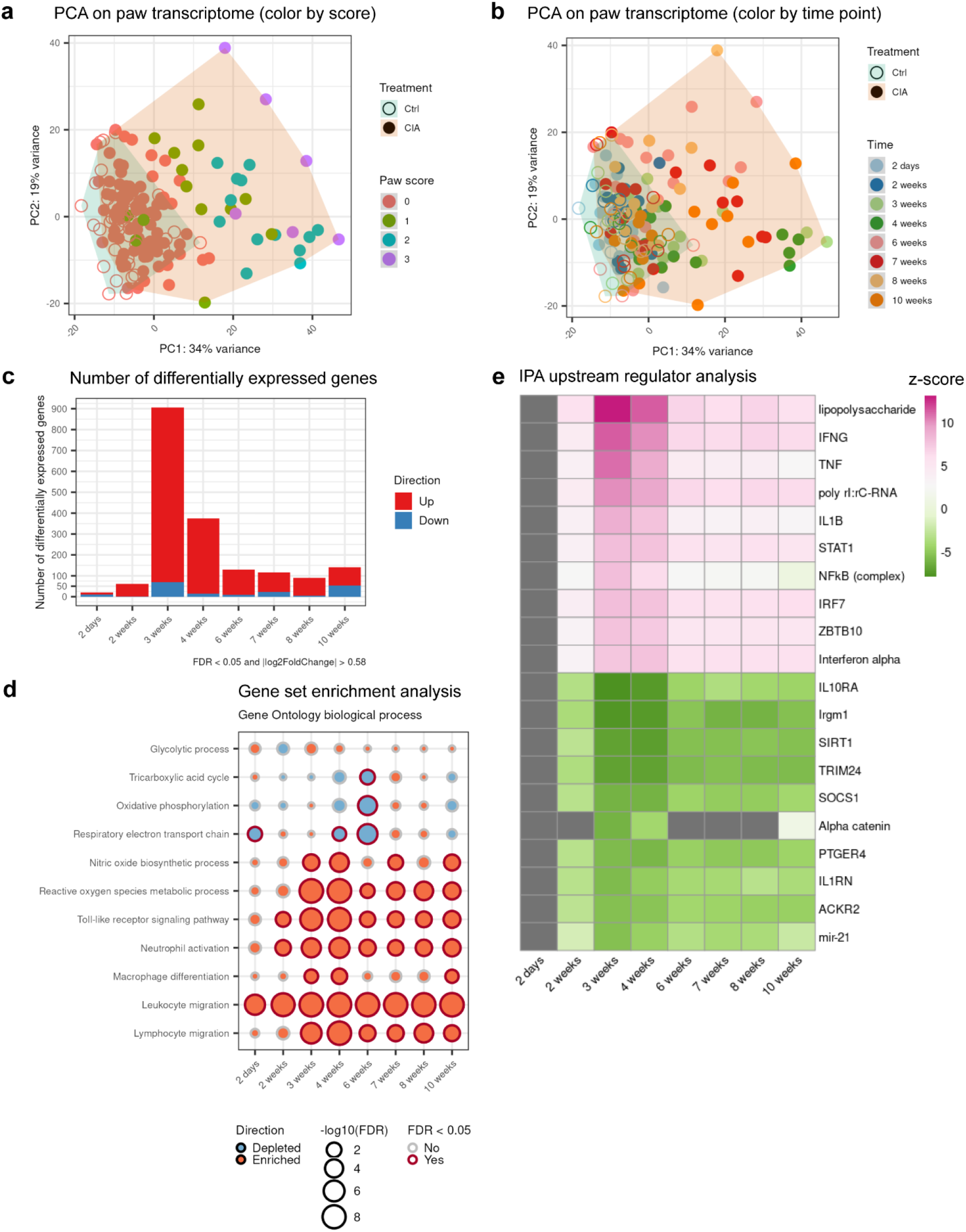
Transcriptional profiling of CIA paws over disease progression revealed innate immune cell activation and metabolic adaptations. Bulk RNA-seq on right hind paws from control and CIA mice at different disease stages. At each time point, the sample size for controls is ≥5 and the sample size for CIA is ≥ 14. **a** PCA score plot with paw samples color-coded by their respective arthritis scores. **b** PCA score plot with paw samples color-coded by their respective sacrifice time points. **c** Number of differentially expressed genes at each time point (CIA samples compared to the controls). The likelihood ratio test (LRT) in DESeq2 was performed for statistical analysis. Significance was decided by cutoffs of FDR < 0.05 and fold change > 1.5 in up or down direction. **d** Enriched or depleted Gene Ontology biological process gene sets at each time point. Gene set enrichment analysis was carried out using the R package fgsea. **e** Top 10 activated and top 10 inhibited upstream regulators from analysis by IPA. Only upstream regulators with Benjamini-Hochberg (B-H) adjusted p-values < 0.05 were shown with z-scores on a color gradient (activated: pink, inhibited: green). For upstream regulators with B-H adjusted p-values ≥ 0.05, the z-scores were shown as grey.

To decipher the molecular mechanism in different stages of arthritis development, we performed gene set enrichment analysis against the biological process gene sets from Gene Ontology database [26] (Fig. 3d). The gene sets of TCA cycle, OXPHOS and respiratory electron transport chain were depleted in CIA samples. In line with the knowledge that disruption in OXPHOS can generate reactive oxygen species (ROS) [27], gene sets of nitric oxide biosynthetic process and ROS metabolic process were enriched in CIA samples. These changes in oxidative stress pathways aligned with the observed changes in gamma-glutamyl amino acids and NAD+ metabolism (Fig. 2f). We also identified at the early disease time point a strong enrichment of immune-related pathways, namely innate immunity, including toll-like receptors signaling pathway, neutrophil activation, macrophage differentiation and leukocyte migration (Fig. 3d) pointing towards a partial parallel co-expression of immune-and metabolic pathways in the paws.

Next, we wondered which targets induced the observed metabolic and innate immunity pathways. Upstream regulator analysis using Ingenuity Pathway Analysis (IPA) showed that lipopolysaccharide (LPS), Interferon gamma (IFNG), TNF and Nuclear factor kappa B (NFκB) were predicted to be activated from week two onwards, which is consistent with the enrichment of the gene sets of innate immunity (Fig. 3d, e). On the other hand, immunity-related GTPase family M member 1 (Irgm1) and Sirtuin 1 (SIRT1) were predicted to be inhibited regulators. Fibroblasts lacking Irgm1 were reported to have dysfunctional mitochondria and increased ROS [28]. SIRT1 is an energy sensor and a histone deacetylase that broadly regulates cellular metabolism. SIRT1 deacetylates various transcription factors such as PPARγ co-activator 1 alpha (PPARGC1A), NFkB and HIF1A and thereby plays a critical role in mitochondrial biogenesis, cellular metabolism, protection against oxidative stress and inflammation [29, 30]. These findings are in line with the results of our gene set enrichment analysis, where OXPHOS is depleted and ROS production is enriched. Furthermore, SIRT1 uses NAD+ as a co-substrate, generating nicotinamide as a by-product [22]. Therefore, inhibition of SIRT1 in upstream regulator analysis is also consistent with reduced level of nicotinamide in plasma.

### Tissue MALDI-MS imaging confirmed reduction of NAD+ in CIA paws

We then used MALDI-MS imaging to monitor metabolite distribution on tissue sections of mouse paws. The imaging methodology caught 11514 unique m/z signals. PCA showed a clear separation between controls and CIA samples (Fig. 4a). Within the CIA samples, the paws with higher arthritis severity (score = 2 or 3) tended to cluster on the left part of the PCA plot. In addition, we observed a partial separation between CIA samples of different disease time points (Fig. 4b). We then focused on the signal distribution of NAD+, the co-substrate of SIRT1, on the paw sections (Fig. 4c). Quantifying the NAD+ signal indicated that its level was downregulated in CIA paws at week two, four and six (Fig. 4d). This supports the findings of SIRT1 inhibition in CIA paws (Fig. 3e) and depletion of nicotinamide in CIA plasma (Fig. 2g).

**Fig. 4:**
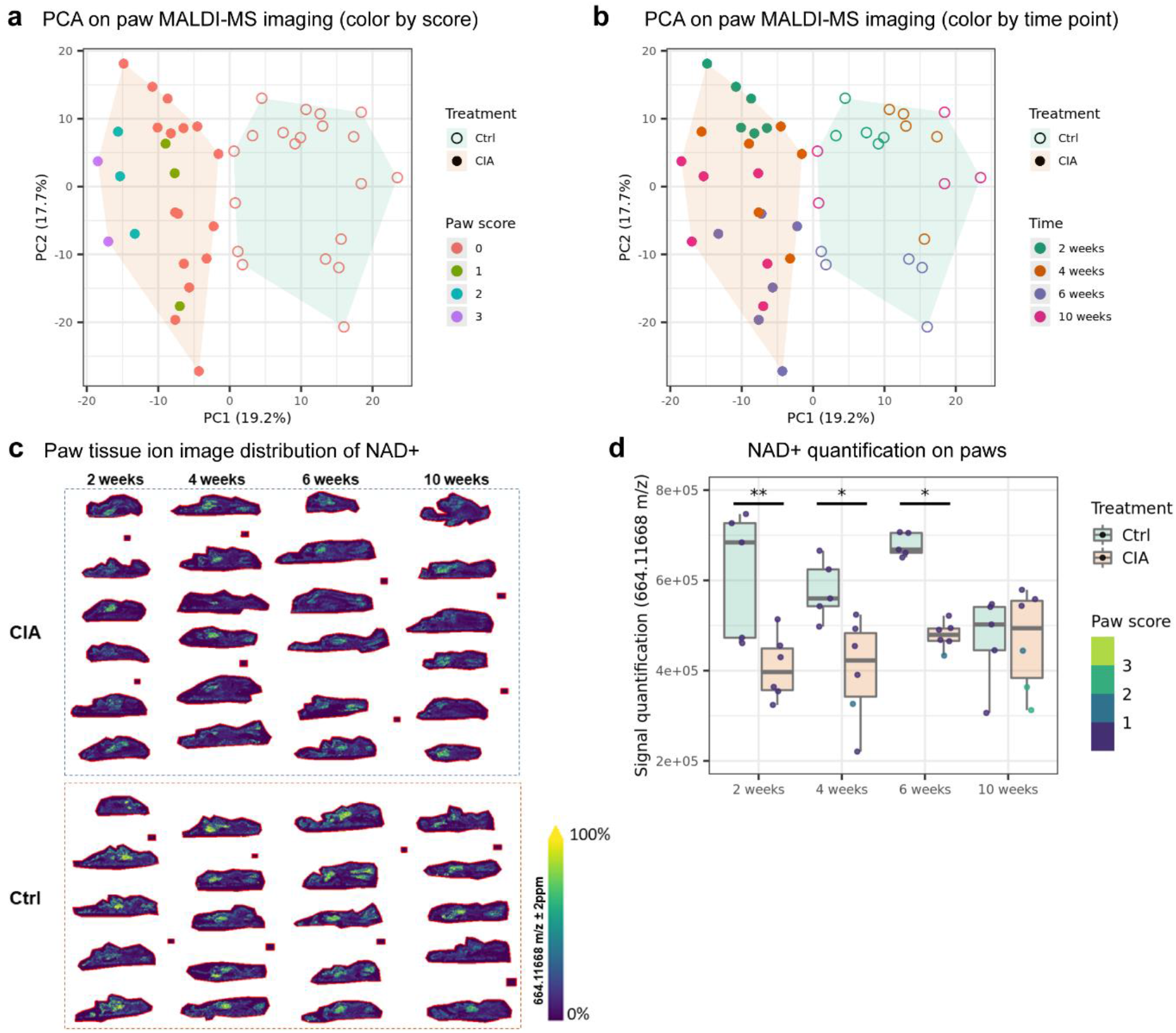
Tissue MALDI-MS imaging showed reduction of NAD+ in CIA paws. Tissue MALDI-MS imaging on left front paws from control and CIA mice at different disease stages. At each time point, the sample size for controls is 5 and the sample size for CIA is 6. **a** PCA score plot with paw samples color-coded by their respective arthritis scores. **b** PCA score plot with paw samples color-coded by their respective sacrifice time points. **c** NAD+ distribution on paw tissue ion image per mouse. Cryosections of mouse paws were analyzed by MALDI-MS tissue imaging mass spectrometry. The mass to charge 664.11668 was assigned to NAD+ with a mass accuracy of 0.4 ppm and its TIC (T=Total Ion Count) normalized tissue distribution is shown at a spatial lateral resolution of 50×50 μm. The color legend reflects the relative intensity [Arb.U] distribution from 0 to 100 %. **d** Image signal quantification of NAD+ on paws. Two-way ANOVA with Tukey multiple pairwise-comparisons was done (p adj < 0.05 *, p adj < 0.01 **, p adj < 0.001 ***).

### Multis-omics factor analysis identified factors correlating with arthritis severity

Combined analysis of omics datasets has dramatically increased both, our understanding and the validation of pathways and targets. Therefore, we assembled the transcriptome data from paws and the metabolomics data from plasma from in total 171 overlapping mice and run multi-omics factor analysis (MOFA). MOFA is an unsupervised method which infers a set of low-dimensional data representations, factors, which drive the major variations among the data [31] (Fig. 5a). MOFA identified 6 factors (Fig. 5b), based on the criterion that a factor should account for > 2% of the variance in at least one omics dataset. Among these, Factors 1 and 2 were mainly active in the transcriptome data. In contrast, Factor 3 was mainly specific to metabolomics data. Cumulatively, the 6 factors explained nearly 20% of variation in metabolomics data and 70% of transcriptome data (Fig. S1).

**Fig. 5:**
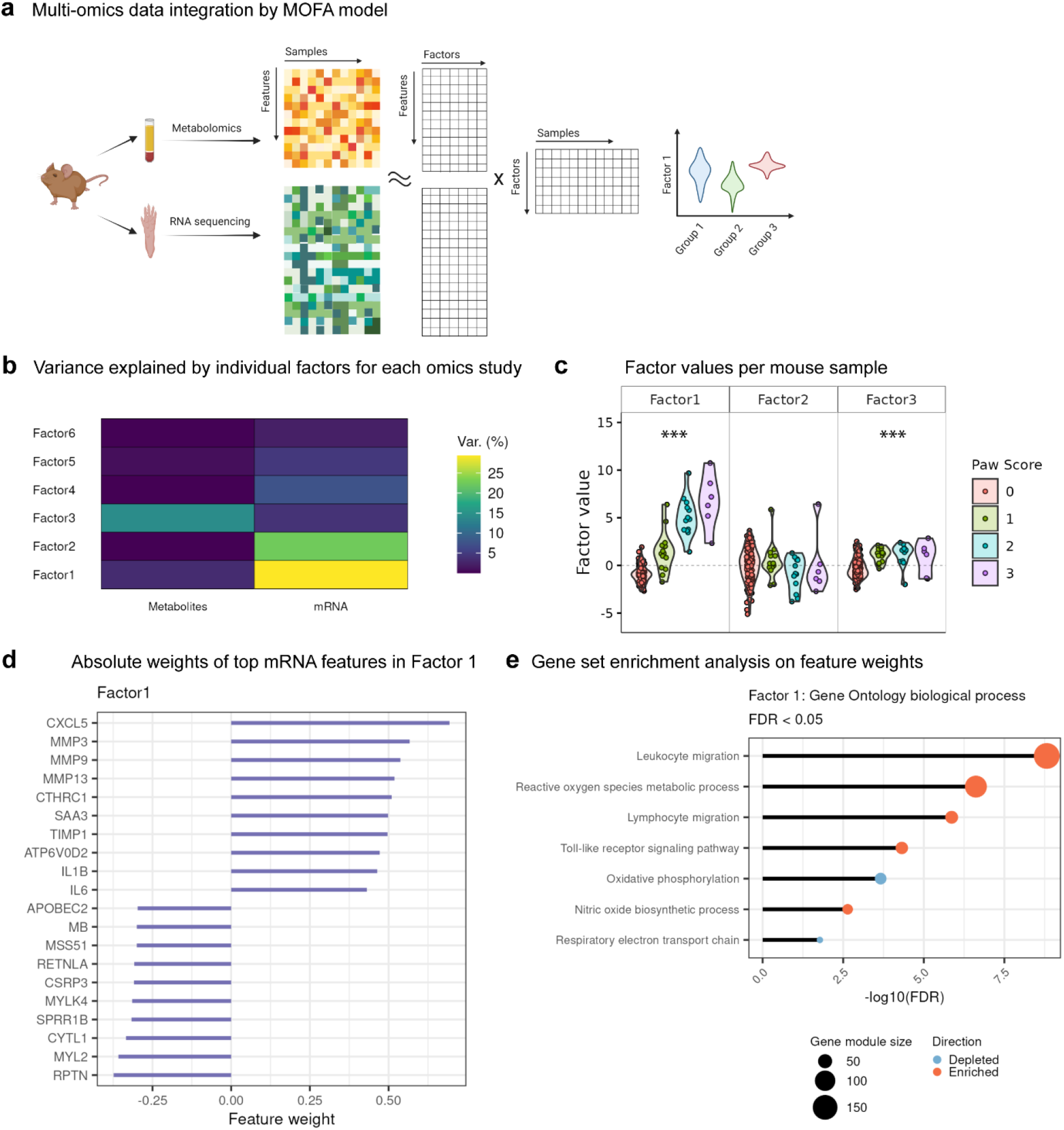
Multi-omics data integration identifies ROS as disease-correlated factor. **a** Overview of MOFA model. MOFA [31] is an unsupervised method for integrating multi-omics data. In this study, MOFA decomposes the data matrices of metabolomics and transcriptomics with co-occurrent samples into a weight matrix for each omics data and a matrix of factors for each sample. “Features” denote metabolites in metabolomics data, or mRNA in transcriptome data. Visualization of sample groups by factor values can identify factors associated with disease phenotypes. **b** Proportion of total variance (Var.) explained by individual factors for each omics experiment. **c** Visualization of samples using Factors 1, 2 and 3. The dots on violin plots represent mouse subjects. The colors denote the arthritis scores of the right hind paws. One-way ANOVA was used to assess if there is any significant difference among the subjects in the 4 score groups (p < 0.05 *, p < 0.01 **, p < 0.001 ***). **d** Absolute weights of the top features of Factors 1 in transcriptome data. **e** Gene set enrichment analysis on the feature weights of mRNA in Factor 1 (FDR < 0.05).

We wondered whether these factors would be able to describe the different stages of disease or disease scores. Factor 1 clearly separated the four groups of paw arthritis scores, which suggests that Factor 1 is able to represent the arthritis severity of mice (Fig. 5c). To investigate the aetiology of Factor 1, we plotted the weights of the top mRNA features on Factor 1 (Fig. 5d). The neutrophil-recruiting chemokine Cxcl5 [32] and the joint-degrading matrix metalloproteinases (MMPs) [33] were assigned the highest positive weights in Factor 1, suggesting that Factor 1 is positively associated with neutrophil activation and joint damage. To further understand the molecular functions of Factor 1, we performed gene set enrichment analysis on the feature weights of mRNA in Factor 1 (Fig. 5e). We found enrichment in the gene sets of ROS production and toll-like receptor signaling pathways, and depletion in the gene sets of OXPHOS and respiratory electron transport chain, which are in line with the analysis results on bulk RNA-seq of mouse paws. Taken together, MOFA results suggest that Factor 1 serves a representation of arthritis severity in mice and is related to ROS production and disruption in mitochondrial respiration.

### Pro-inflammatory macrophage transcriptome changes partially overlapped with early CIA transcriptome changes

The highest elevated energy metabolite in CIA mouse plasma was itaconate (Fig. 2g), a signature metabolite of pro-inflammatory macrophage activation [25]. In addition, we observed significant upregulations in several innate immunity pathways and innate upstream regulators like LPS (Fig. 3d, e). We therefore hypothesized that a large proportion of metabolic programming in arthritis inflammation could be driven by macrophages. We next investigated the transcriptome by bulk RNA-seq on human primary monocyte-derived macrophages after LPS stimulation. PCA on macrophage transcriptome showed a clear separation between LPS-stimulated and unstimulated macrophages (Fig. 6a). We then compared the significant differentially expressed genes between CIA paws at week three and LPS-stimulated human macrophages (Fig. 6b). Although the overall overlap was limited due to the drastic changes on macrophages stimulated in an *in vitro* setting, we found 254 shared upregulated genes, which reflects ~30% (254/838) of significantly differentially upregulated genes in CIA paws at week three.

**Fig. 6:**
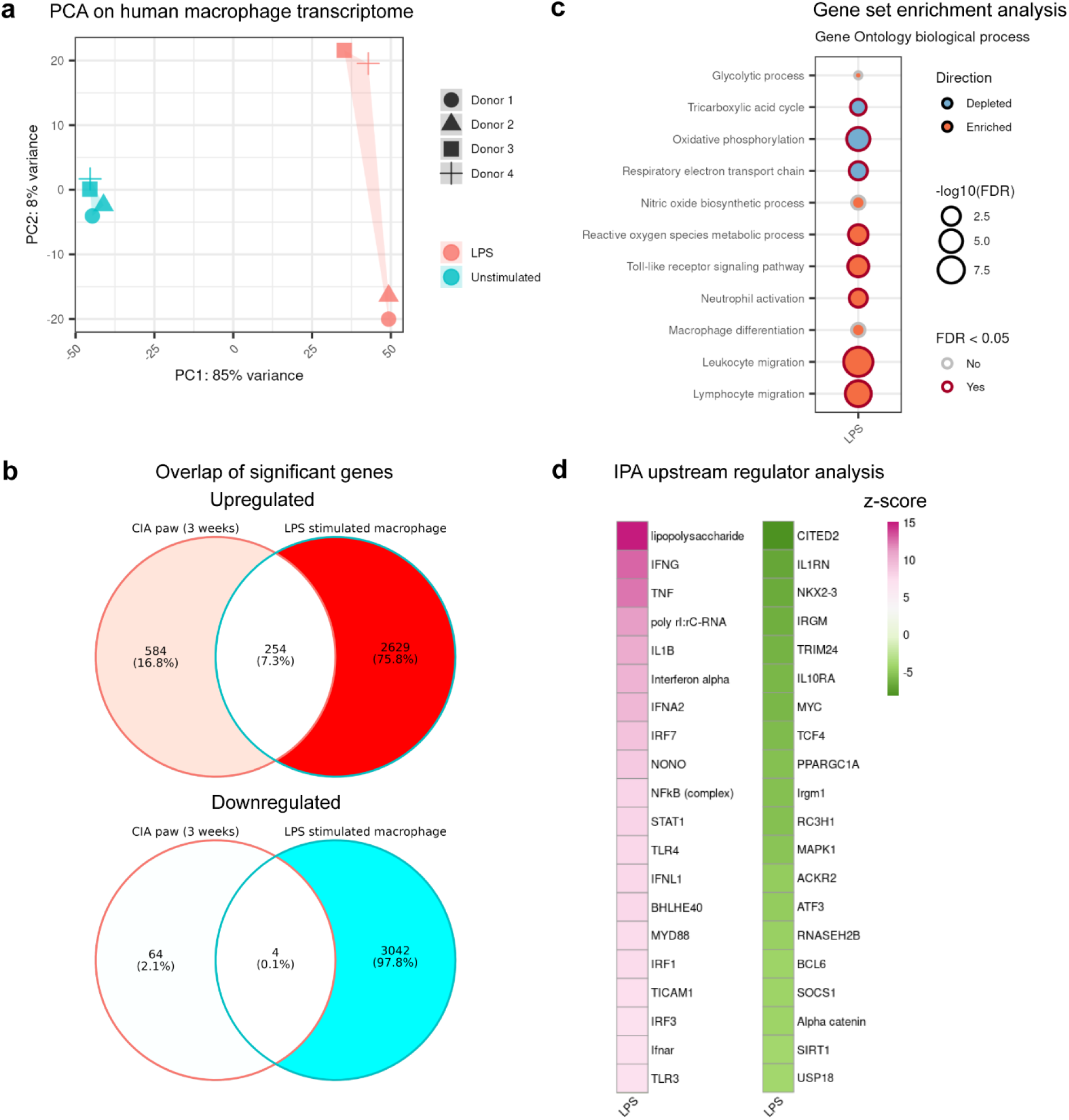
Transcriptional profiling of LPS-stimulated human macrophages partially overlaps to early transcript changes in mouse CIA. Bulk RNA-seq analysis of human monocyte-derived macrophages from four human donors. Stimulation was done with 50 ng/ml LPS for 24 hours. **a** PCA score plot with four samples of human macrophages before and after LPS stimulation. **b** Overlap of significant differentially expressed genes between CIA paws at week three and LPS-stimulated macrophages. The Wald test in DESeq2 was performed for statistical analysis on macrophage dataset and LPS-stimulated samples were compared to the unstimulated. **c** Enriched or depleted Gene Ontology biological process gene sets. **d** Top 20 activated and top 20 inhibited upstream regulators from analysis by IPA. Only upstream regulators with Benjamini-Hochberg (B-H) adjusted p-values < 0.05 were shown with z-scores on a color gradient (activated: pink, inhibited: green).

Gene set enrichment analysis on LPS-stimulated human macrophage transcriptome also yielded similar results as the analysis on CIA transcriptome (Fig. 6c). The gene sets of OXPHOS and respiratory electron transport chain were depleted upon LPS stimulation, whereas gene sets of ROS metabolism, toll-like receptor signaling pathway and innate immunity were enriched. Toll-like receptor 4 (TLR4) is required for LPS recognition and TLR4 signaling leads to accumulation of HIF1A [34]. Previous studies have linked activation of HIF1A closely to ROS generation and increase of glycolytic activity [35]. Upstream regulator analysis using IPA on LPS-stimulated macrophages was also generally in line with the analysis on CIA paws, where LPS, IFNG, TNF and NFκB were predicted to be activated (Fig. 6d). Furthermore, SIRT1 and Irgm1 (Fig. 6d) were predicted to be inhibited. Overall, we observed consistent metabolic and immunological changes on transcriptome level between CIA paws and LPS-stimulated macrophages.

### Metabolic dysregulation confirmed by transcriptome changes in various stages of RA patient disease progression

To validate the findings on CIA paws in a context of human RA, we looked into a published RNA-seq dataset (GSE89408) [36] on synovium biopsies from healthy humans (n=27) and patients with arthralgia (n=10), undifferentiated arthritis (UA, n=6) and early RA (n=57). The pathogenesis of human RA (Fig. 7a) originates from asymptomatic autoimmunity, which manifests into arthralgia with nonspecific joint symptoms and further progresses to UA with clinical synovitis. Subsequently UA advances into early stage of RA, which is defined as within 12 months of diagnosis without prior treatment. PCA on human synovium transcriptome revealed clear separation between the healthy group and the clinically symptomatic groups (Fig. 7b). In contrast, the early RA group overlapped with the UA group and partially overlapped with the arthralgia group in PCA. This suggests that although there is a clear divergence between the healthy and the diseased, much less differentiation exists between the different disease stages. We performed the differential gene expression test (Fig. 7c) and then compared the significant differentially upregulated genes between CIA paws at week three and human synovium (Fig. 7d). We found 310 shared upregulated genes between CIA paws at week three and early RA human synovium, which reflects ~37% (310/838) of significant upregulated genes in CIA paws at week three. In addition, the overlap was most pronounced with early RA (Fig. 7d). It should be mentioned though, that the human RA study revealed a huge amount of transcript changes which are only partially seen in mouse CIA (Fig. 7d).

**Fig. 7:**
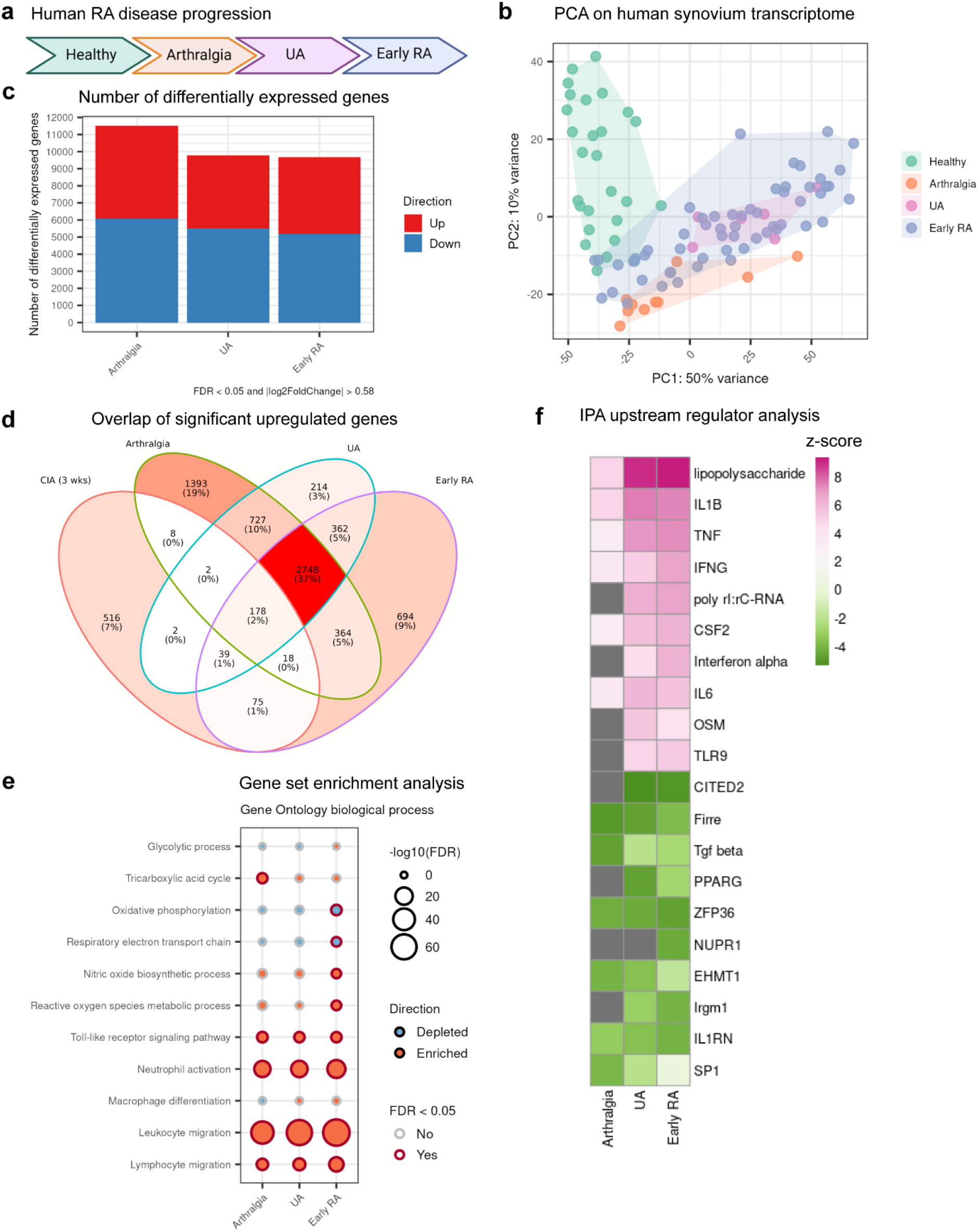
Gene expression of human synovium over disease progression to RA showed similar changes in oxidative stress and innate immunity. Published RNA-seq dataset (GSE89408) contains synovium samples from healthy humans (n=27) and patients with arthralgia (n=10), undifferentiated arthritis (UA, n=6) and early RA (n=57). **a** Human RA progression from arthralgia, UA and early RA. **b** PCA score plot with patient samples color-coded by disease stage. **c** Number of significant differentially expressed genes at each disease stage. The Wald test in DESeq2 was performed for statistical analysis and the diseased samples (arthralgia, UA and early RA) were compared to the healthy. Significance was decided by cutoffs of FDR < 0.05 and fold change > 1.5 in up or down direction. **d** Overlap of significant differentially upregulated genes between CIA paws at week three and human synovium of different RA stages. **e** Enriched or depleted Gene Ontology biological process gene sets. **f** Top 10 activated and top 10 inhibited upstream regulators from analysis by IPA. Only upstream regulators with Benjamini-Hochberg (B-H) adjusted p-values < 0.05 were shown with z-scores on a color gradient (activated: pink, inhibited: green). For upstream regulators with B-H adjusted p-values ≥ 0.05, the z-scores were shown as grey.

Despite these limitations, gene set enrichment analysis on human synovium transcriptome was also largely in line with the analysis on CIA transcriptome (Fig. 7e). The gene sets of OXPHOS and respiratory electron transport chain were depleted while the gene sets of ROS metabolism enriched. In addition, we also observed an enrichment of gene sets of innate immunity, including toll-like receptor signaling pathway, neutrophil activation, and leukocyte migration. Furthermore, upstream regulator analysis using IPA on human synovium biopsies gave rise to similar predictions as the analysis on CIA paws (Fig. 7f, Supplementary Table 2), although SIRT1 signaling was - despite being significant- to a lower extent inhibited. Comparing the *in vitro* macrophage, human RA and mouse CIA studies, we identified the following consistent upstream regulators: LPS, IFNG, TNF and NFκB were upregulated, while Irgm1 and SIRT1 were inhibited across the studies. Overall, we were able to confirm and translate the metabolic reprogramming in CIA paws with the early human RA synovium RNA-seq results.

### Consistent changes in oxidative stress, glycolysis and NFkB targets

To query the metabolic reprogramming at individual gene level, we looked into the changes of major metabolic and innate immunity genes in the significantly altered pathways identified via the gene set enrichment analysis. NFκB mediated pro-inflammatory molecules in the immune activation, such as IL6 and IL1B, showed consistent increase of gene expression (Fig. 8a). In line with the increased ROS production, we observed significant upregulation of myeloperoxidase (MPO) and NADPH oxidase subunits (NCF1, NCF2 and NCF4) in CIA paws (Fig. 8a). Differential gene expression on LPS-stimulated human macrophages and human RA synovium biopsies partially confirmed this finding (Fig. 8b-c). MPO is involved in ROS generation [37] and is required for neutrophil extracellular trap (NET) formation [38]. NETs have been implicated in microbial defense and served as a source of autoantigens in RA [39]. In addition, NADPH oxidase is critical for ROS production in neutrophils and macrophages [40].

**Fig. 8:**
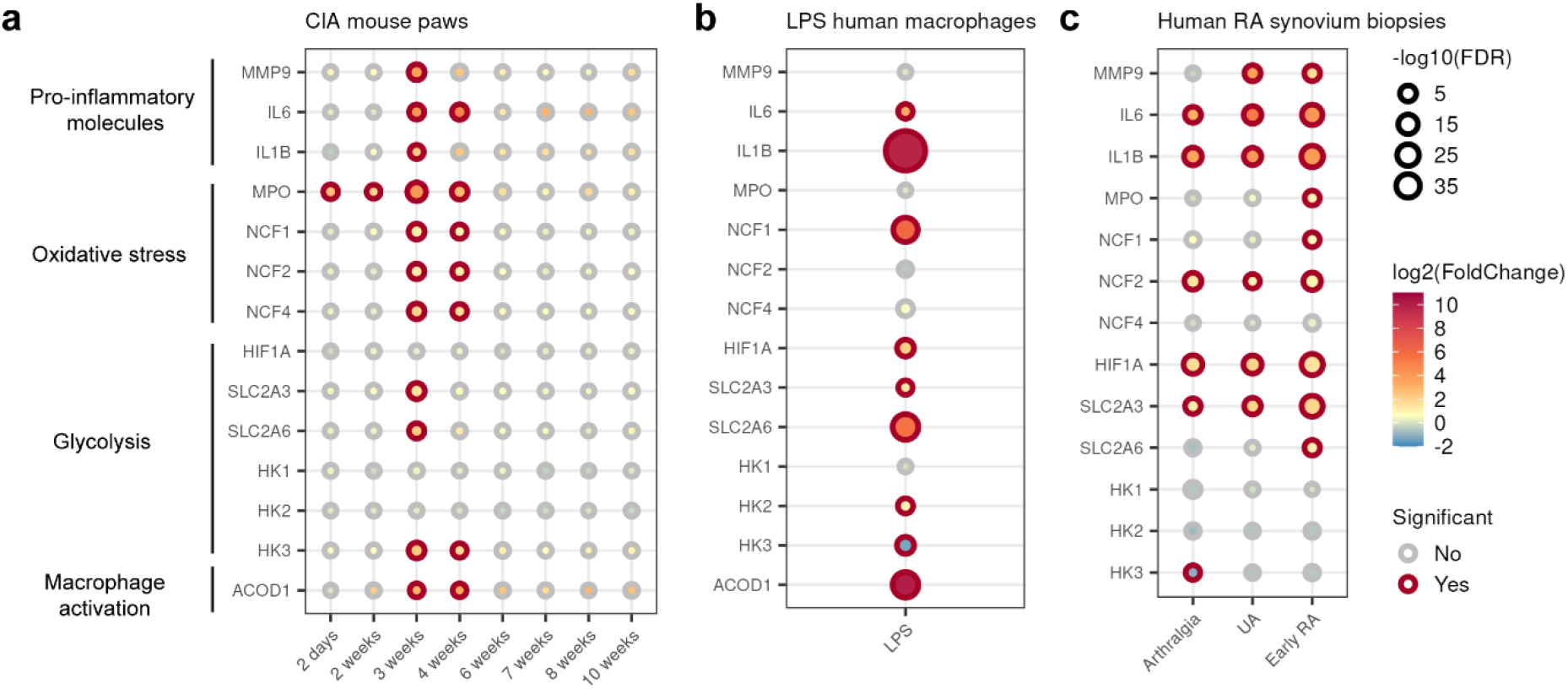
Metabolic reprogramming in early RA. **a-c** Differential gene expression of major metabolic genes in the RNA-seq studies of CIA mouse paws, LPS-stimulated human macrophages, and human RA synovium biopsies. Significance was decided by cutoffs of FDR < 0.05 and fold change > 1.5 in up or down direction.

The NAD-dependent SIRT1 also directly regulates cellular metabolism via inhibiting the key transcription factor HIF1A by deacetylation [41]. HIF1A, was significantly upregulated in LPS-stimulated human macrophages and synovium biopsies of RA patients (Fig. 8b-c). HIF1A is one of the mediators of SIRT1 regulating immunometabolism by promoting expression of genes involved in glycolysis [42]. Upon examining the differential expression of glycolytic genes, we observed consistent upregulation in glucose transporters GLUT3 (SLC2A3) and GLUT6 (SLC2A6, Fig. 8a-c).

Increased glycolysis and a broken TCA cycle are characteristics of pro-inflammatory macrophages. ACOD1 encodes an itaconate-producing enzyme, shifting TCA cycle towards itaconate production in pro-inflammatory macrophages [43]. ACOD1, was also significantly upregulated in CIA paws and LPS-stimulated human macrophages (Fig. 8a-b). ACOD1 expression in human synovium RNA-seq dataset was too low and therefore it was filtered out before differential gene expression analysis (Fig. 8c).

## Discussion

Understanding the pathogenesis of RA is essential for development of new therapeutics. However, few studies have systemically examined the early metabolic changes in arthritis. Here, we utilized RNA-seq and metabolomics to provide a time-resolved comprehensive overview on the arthritis progression in CIA mice. We then used an unsupervised multi-omics factor analysis method to identify the important metabolic processes in etiology of RA development. Furthermore, we validated the major findings of the CIA study with two human datasets, including LPS-stimulated macrophages and synovium biopsies of RA patients.

The onset of clinical RA is preceded by a phase of asymptomatic autoimmunity where innate immunity plays a key role in disease progression [2, 44]. In human RA synovium, neutrophils and macrophages are the highly abundant cell types [45–47] and the number of synovial macrophages may predict joint destruction [48]. Furthermore, activation of innate immune cells was also observed in synovium biopsies from treatment-naïve early RA [3]. In our study, we observed a substantial increase of the pro-inflammatory macrophage activation marker itaconate in plasma and neutrophil activation marker MPO in the paws, just two days after the first immunization. These early innate changes were confirmed by both gene set enrichment and upstream regulator analysis. In addition, 30-37% of the CIA gene differential expression could be explained by the transcriptome changes of LPS-stimulated human macrophages and human synovium of early RA. We therefore hypothesize that innate immune cells have a critical role in the early pathology in CIA paws.

The observed metabolic changes were also in alignment with innate immune cell activation and infiltration. We report that metabolic changes at both gene expression and metabolite levels occurred already at the asymptomatic stages of the CIA mouse model (day two and week two). There, gamma-glutamyl amino acids were significantly increased and the NAD+ precursor in salvage pathway, nicotinamide, was depleted in CIA plasma. In addition, MALDI-MS imaging confirmed the depletion of NAD+ in CIA paws. Interestingly, in line with our findings, the metabolite of nicotinamide, 1-methylnicotinamide, was significantly decreased in plasma of active RA patients [49]. Furthermore, plasma of RA patients showed dysregulated glycolysis, TCA and amino acid metabolites [49]. Our identification of itaconate was also reported in the serum/urine of Tg197 arthritis mouse model [50]. Recently, itaconate was identified in an unbiased metabolomics study as early biomarker in RA [51]. This highlights the translational relevance of our longitudinal study and hopefully encourages the validation of some novel described metabolites like gamma-glutamyl amino acids in the future.

Direct comparison of mouse and human datasets in the current study unveiled a shared pattern of enhanced glycolysis, depleted OXPHOS and increased ROS synthesis in arthritis development. Glycolysis provides activated cells with ATP and precursors for biosynthesis via influx into PPP [5]. Inhibition of glycolysis can reduce disease severity of RA mouse models [52, 53]. The upstream regulator analyses on all three transcriptome datasets (Fig. 3e, 6d, 7f) predicted that the key metabolic sensor, SIRT1, was inhibited, while its downstream target NFξB and pro-inflammatory cytokines were activated. This is consistent with the experimental evidence on SIRT1-knockout macrophages, which exhibit hyperacetylated NFkB, higher NFkB-mediated gene transcription and subsequently increased pro-inflammatory cytokine production [30]. Similarly, treatment of RA macrophages with resveratrol, an unspecific SIRT1 activator, enhanced anti-inflammatory M2 and blocked pro-inflammatory M1 polarization [54]. These authors confirmed their findings with SIRT1 knockdown in RA macrophages and in bone-marrow derived macrophages from SIRT1 transgenic mice, strengthening a direct role of SIRT1 in macrophage polarization. These mice also showed improved arthritis scores in a CIA model accompanied by an increase in M2 macrophage markers. Similarly, resveratrol improved arthritis scores and reduced incidence in the CIA model - both in preventive and therapeutic mode - by inhibiting Th17 and B cell function [55]. Furthermore, a recent study also identified gene polymorphisms of SIRT1 in RA patients [56].

NAD+ integrates the mRNA and metabolite changes due to its cofactor role for SIRT1’s enzymatic activity [22]. We hypothesize that at the start of autoimmunity in RA, macrophages and neutrophils experience a change of their effector functions in the direction of pro-inflammatory activation and tissue destruction (Fig. 9). The activation of these innate immune cells is accompanied by increased glycolysis that consumes NAD+, decreased OXPHOS that slows regeneration of NAD+ by Complex I, increased ROS production and increased intracellular availability of amino acids. Low NAD+ level causes inhibition of SIRT1, which results in accumulation of HIF1A and activation of NFkB. HIF1A and NFkB pathways further promote glycolysis over OXPHOS, which serves as a positive feedback loop in the manifestation of inflammation. On top of that, our data is in line with the notion that macrophages and neutrophils contribute to synovitis by ROS production in MOFA analysis. Our result further underlines the role of SIRT1 and macrophage metabolism in the development of arthritis.

**Fig. 9:**
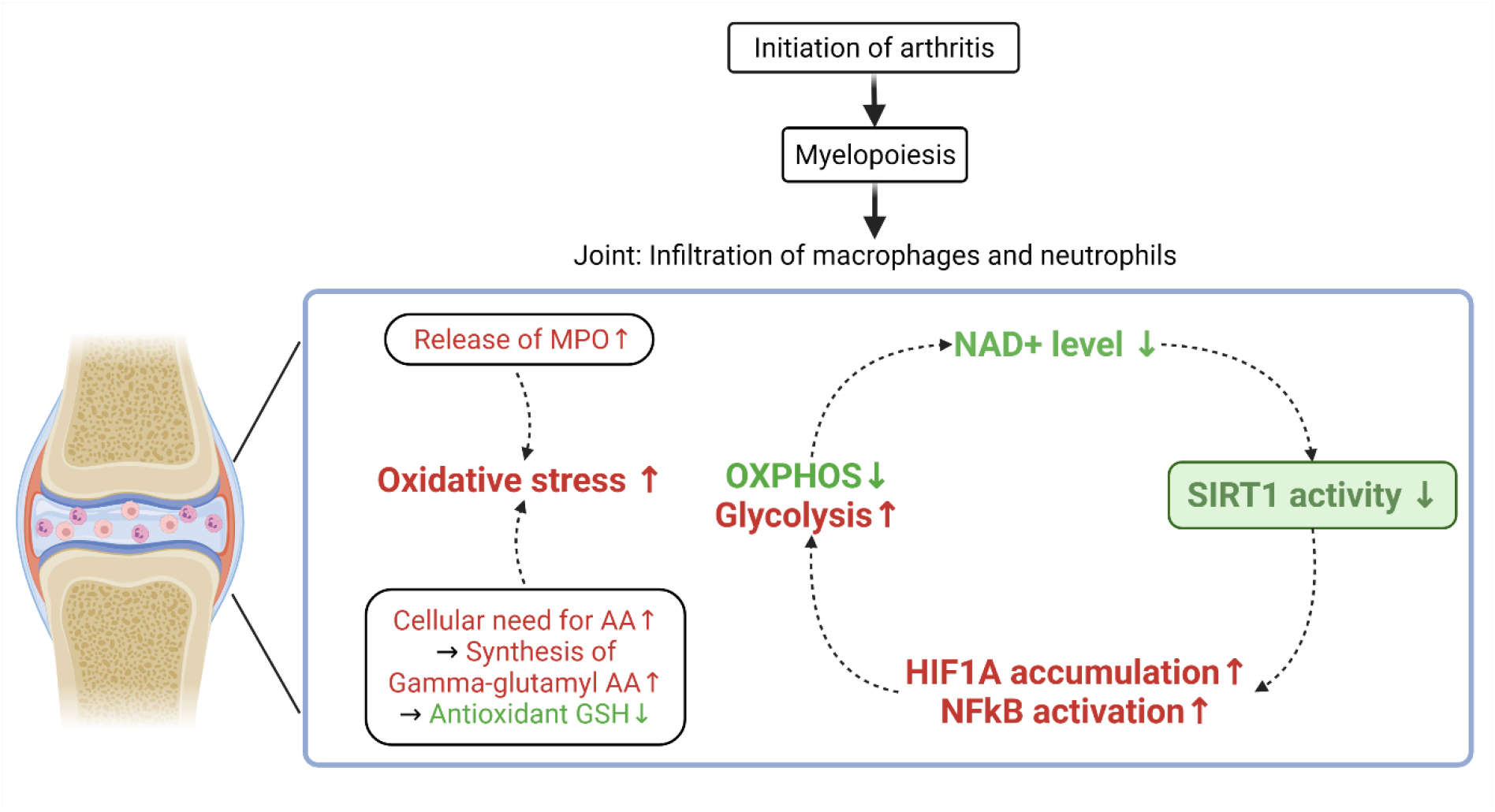
Innate immune cells drive onset of inflammation via SIRT1 axis in CIA mouse model. Schematic showing the metabolic changes driven by SIRT1 axis at the initial phase of RA.

This study also provides a resource that can be mined for mechanistic investigation of metabolic dysregulation that underlies arthritis development in CIA mouse model. We took the large heterogeneity in arthritis phenotype of these mice into consideration and used a large sample number for the CIA mice. Although validation of the CIA RNA-seq study findings with two human datasets does not yield complete match in terms of magnitude of gene expression changes, this is largely because of the differences in the species, tissue types and sequencing protocols. Also, we acknowledge that single-cell RNA-seq on mouse synovium would have provided more comprehensive gene expression at individual cell level, but this approach would have been technically challenging to apply to the tiny size of mouse synovium.

In conclusion, our dynamic analysis of molecular changes on CIA mouse model summarizes the metabolic changes during the development of arthritis. We speculate that activation of SIRT1 and thereby modulation of glycolysis, OXPHOS and ROS metabolism would yield beneficial effect in treatment of RA. We also delivered a comprehensive time-resolved plasma metabolome and paw transcriptome for further pursuit of causal implication in immunometabolism.

## Methods

### Collagen-induced arthritis (CIA) in mice

All animal experiments were performed in compliance with German animal welfare legislation implementing Directive 63/2010/EU, approved by the Regional Council (Regierungspräsidium Darmstadt) of the state of Hessen, Darmstadt, Germany and carried out in an Association for Assessment and Accreditation of Laboratory Animal Care (AAALAC) International accredited facility.

Chicken type II collagen (CII, MD Bioproducts) was dissolved overnight in 0.1 M acetic acid (Fulka) at 4°C. The solution was then emulsified with an equal volume of Complete Freund’s adjuvant (CFA, Sigma) in an ice-cold water bath. The male DBA1 mice (age between 10 to 12 weeks, ENVIGO) were intradermally injected with 50 μl of this emulsion (containing 100 μg CII/mouse) at the base of the tail (day 0). On day 21 animals were boosted with 50 μl emulsion of CII in Incomplete Freund’s adjuvant (IFA, Sigma). The arthritis severity in CIA mice was scored in all four paws at 3 times per week for in total 10 weeks. The severity was scored as follows: 0 – no clinical sign of arthritis; 1 – visible swelling limited to wrists and ankles; 2 – visible swelling expanded to the pads with slight redness; 3 – visible swelling with heavy redness and ankylosis.

Tissue collections were done at 9 different time points during the experiment, where age matched male DBA1 mice (controls) were sacrificed at the same time as the CIA mice. At sacrifice, the spleen was weighted fresh. Blood was collected into EDTA-coated micro sample tubes (Sarstedt) after cardiac puncture. Plasma was extracted after centrifugation at 1,530 x g for 15 min at 4°C and stored at −80°C. Mouse paws were cut and snap frozen in liquid nitrogen.

### Human macrophage differentiation and stimulation

Human whole blood was from the Sanofi in-house blood donor service. The protocol was approved by the local ethics committee and all blood donors signed informed consent. Human peripheral blood monocytes were isolated from anonymous donors’ prefinal blood using Ficoll density centrifugation, followed by magnetic separation with positive selection (CD14 MicroBeads, Miltenyi Biotec). Monocytes were differentiated into macrophages by culturing in macrophage serum-free medium (Life Technologies) containing 50 ng/ml recombinant human macrophage colony-stimulating factor (Immunotools) for 5 days. Following differentiation, cells were cultured in RPMI 1640 medium supplemented with 10% heat inactivated fetal bovine serum (Thermo Fisher) and maintained at 37°C in a 5% CO_2_/air environment. Macrophages were stimulated with 50 ng/ml lipopolysaccharides from Escherichia coli O111:B4 (Sigma) for 24 hours.

### Luminex assays

Mouse magnetic Luminex assays (R&D Systems) were performed on EDTA mouse plasma according to the manufacturer’s instructions. Beads were read on a Luminex MAGPIX Instrument using the xPONENT software (Luminex Corporation).

### Untargeted metabolomics with ultrahigh performance liquid chromatography-tandem mass spectroscopy (UPLC-MS/MS) on mouse plasma

Untargeted metabolomics on mouse plasma was performed at Metabolon Inc. (Morrisville, NC, USA). Samples were prepared using the automated MicroLab STAR system from Hamilton Company. Several recovery standards were added prior to the first step in the extraction process for QC purposes. Proteins were precipitated with methanol under vigorous shaking for 2 min (Glen Mills GenoGrinder 2000) followed by centrifugation. The resulting extract was divided into five fractions: two for analysis by two separate reverse phase (RP)/UPLC-MS/MS methods with positive ion mode electrospray ionization (ESI), one for analysis by RP/UPLC-MS/MS with negative ion mode ESI, one for analysis by HILIC/UPLC-MS/MS with negative ion mode ESI, and one sample was reserved for backup. Samples were placed briefly on a TurboVap (Zymark) to remove the organic solvent. The sample extracts were stored overnight under nitrogen before preparation for analysis.

All methods utilized a Waters ACQUITY ultra-performance liquid chromatography (UPLC) and a Thermo Scientific Q-Exactive high resolution/accurate mass spectrometer interfaced with a heated electrospray ionization (HESI-II) source and Orbitrap mass analyzer operated at 35,000 mass resolution. The sample extract was dried then reconstituted in solvents compatible to each of the four methods. Each reconstitution solvent contained a series of standards at fixed concentrations to ensure injection and chromatographic consistency. One aliquot was analyzed using acidic positive ion conditions, chromatographically optimized for more hydrophilic compounds. In this method, the extract was gradient eluted from a C18 column (Waters UPLC BEH C18-2.1×100 mm, 1.7 μm) using water and methanol, containing 0.05% perfluoropentanoic acid (PFPA) and 0.1% formic acid (FA). Another aliquot was also analyzed using acidic positive ion conditions, however it was chromatographically optimized for more hydrophobic compounds. In this method, the extract was gradient eluted from the same afore mentioned C18 column using methanol, acetonitrile, water, 0.05% PFPA and 0.01% FA and was operated at an overall higher organic content. Another aliquot was analyzed using basic negative ion optimized conditions using a separate dedicated C18 column. The basic extracts were gradient eluted from the column using methanol and water, however with 6.5 mM Ammonium Bicarbonate at pH 8. The fourth aliquot was analyzed via negative ionization following elution from a HILIC column (Waters UPLC BEH Amide 2.1 x 150 mm, 1.7 μm) using a gradient consisting of water and acetonitrile with 10 mM Ammonium Formate, pH 10.8. The MS analysis alternated between MS and data-dependent MSn scans using dynamic exclusion. The scan range varied slighted between methods but covered 70-1000 m/z. Raw data files are archived and extracted as described below.

Several types of controls were analyzed in concert with the experimental samples: a pooled matrix sample generated by taking a small volume of each experimental sample (or alternatively, use of a pool of well-characterized human plasma) served as a technical replicate throughout the data set; extracted water samples served as process blanks; and a cocktail of QC standards that were carefully chosen not to interfere with the measurement of endogenous compounds were spiked into every analyzed sample, allowed instrument performance monitoring and aided chromatographic alignment. Instrument variability was determined by calculating the median relative standard deviation (RSD) for the standards that were added to each sample prior to injection into the mass spectrometers. Overall process variability was determined by calculating the median RSD for all endogenous metabolites (i.e., non-instrument standards) present in 100% of the pooled matrix samples.

### UPLC-MS/MS data extraction and compound identification

Raw data was extracted, peak-identified and QC processed using Metabolon’s hardware and software. Compounds were identified by comparison to library entries of purified standards or recurrent unknown entities. Metabolon maintains a library based on authenticated standards that contains the retention time/index (RI), mass to charge ratio (m/z), and chromatographic data (including MS/MS spectral data) on all molecules present in the library. Furthermore, biochemical identifications are based on three criteria: retention index within a narrow RI window of the proposed identification, accurate mass match to the library +/- 10 ppm, and the MS/MS forward and reverse scores between the experimental data and authentic standards. The MS/MS scores are based on a comparison of the ions present in the experimental spectrum to the ions present in the library spectrum. Additional mass spectral entries have been created for structurally unnamed biochemicals, which have been identified by virtue of their recurrent nature (both chromatographic and mass spectral). Library matches for each compound were checked for each sample and corrected if necessary. This study generated an untargeted metabolomics dataset that comprises a total of 929 biochemicals, 815 compounds of known identity and 114 compounds of unknown structural identity. Peaks were quantified using area-under-the-curve. For studies spanning multiple days, a data normalization step was performed to correct variation resulting from instrument inter-day tuning differences. Essentially, each compound was corrected in run-day blocks by registering the medians to equal one (1.00) and normalizing each data point proportionately.

### Statistical analysis on untargeted metabolomics of mouse plasma

Following log transformation, two-way ANOVA using base R package (v4.0.0) [57] was performed to identify metabolites exhibiting significant differences between treatment groups (CIA vs. controls) and time points. All reported significant results on untargeted metabolomics have adjusted p-value cutoff < 0.05 after using False Discovery Rate (FDR) method to correct for multiple testing. PCA was performed using the corresponding function in MetaboAnalystR 3.0 [58].

### Tissue MALDI-MS imaging (tMSI) on mouse paws

For MALDI-MS imaging experiments 10 μm thick paw cryosections were cut with a cryostat (CM 1950 cryostat, Leica Biosystems), thaw-mounted onto conductive indium thin oxide coated glass slides (ITO, Bruker Daltonik) and subsequently desiccated at room temperature overnight until matrix application or stored at −80°C under vacuum until further use. MALDI matrix application was performed as previously described [59]. Briefly, 2.5 DHB matrix was dissolved at a concentration of 60 mg/ml in ACN/ddH2O/TFA (50/50/0.2, v/v/v). Matrix deposition onto the tissue was performed by spray coating using a SunCollect sprayer (SunChrom), by using 5 cycles in ascending orders (10, 15, 20, 25, 25 μL/min) at a velocity speed of 300 mm/min. Data acquisition was performed on a 7 T Scimax MRMS Fourier transform ion cyclotron resonance (FT-ICR) mass spectrometer (Bruker Daltonik) equipped with a dual ESI/MALDI ion source and a Smartbeam II Nd:YAG Laser (355 nm) under the acquisition control of FTMS Control 2.3.0. Images were acquired in the positive ion mode in the m/z range of 100 to 2,000 using 2Ώ and 2M magnitude mode with a resolving power of ~400,000 at m/z 400 and a free induction decay of 0.8 ms using FlexImaging 5.0 (Bruker Daltonik) by applying 250 laser shots per measurement position at a lateral resolution of 50 μm. To ensure high mass accuracy an internal lock mass calibration was performed on known DHB ion clusters (m/z 273.03936).

For ion image visualization and statical evaluation, raw data were loaded and processed using SCiLS Lab MVS (Version 9.01.12514, Bruker Daltonik). Normalization of signal intensities were performed using the total ion count (TIC) normalization and a global peak assignment was performed at a frequency of 1 % for data reduction. Reduced data were exported into imzml data format using SCiLS Lab and a peak picking based on the SPUTNIK R package [60] was applied to filter spatial relevant m/z information. The spatial filtered m/z list was mapped against Sanofi’s internal biomarker database at a mass accuracy of 1.2 ppm (1.2 ppm reflects mean ± 2 SD of technical variability over the entire mass range). The intensities values of assigned m/z biomarker candidates were exported from SCiLS Lab as Arb.U and used for statistical evaluation.

Two-way ANOVA with Tukey multiple pairwise-comparisons using base R package (v4.0.0) [57] was applied to calculate adjusted p-values. PCA was performed using the corresponding function in MetaboAnalystR 3.0 [58].

### RNA extraction

For mouse paws, frozen tissues were homogenized in 1.2 ml of QIAzol Lysis Reagent (Qiagen) per sample using a TissueLyser (Qiagen) for 2 x 5 min at 30 Hz, 4°C. The lysates were centrifuged for 5 min at 12,000 g, 4°C to remove debris. 250 μl of supernatant was used for RNA isolation using the Direct-zol-96 RNA kit (Zymo Research) by following the manufacturer’s instructions. For human macrophages, frozen cell pellet from each well was lysed by addition of 350 μl of the TRI Reagent solution included in the Direct-zol-96 RNA kit. Total RNA was then isolated from the lysates using the Direct-zol-96 RNA kit. The concentration of RNA samples was measured using the Quant-iT RNA Assay Kit (Invitrogen). The RNA quality (RIN) was assessed by 2100 Bioanalyzer (Agilent Technologies) or 4200 TapeStation (Agilent Technologies).

### Bulk RNA sequencing

For mouse paws, 1 μg of total RNA was used as an input material for library preparation using NEBNext Ultra II Directional RNA Library Prep Kit for Illumina with NEBNext Poly(A) mRNA Magnetic Isolation Module (New England Biolabs). The libraries were multiplexed and paired-end sequenced (2 × 75 bp) on NextSeq 550 instrument (Illumina) using the NextSeq 500/550 High Output Kit (150 cycles). The average sequencing depth is 8.9 million reads per mouse paw sample.

For human macrophages, 60 ng of total RNA was used for library preparation using NEBNext Ultra II Directional RNA Library Prep Kit for Illumina with NEBNext rRNA Depletion Kit (New England Biolabs). The libraries were paired-end sequenced (2 × 75 bp) on NovaSeq 6000 instrument (Illumina) using the NovaSeq 6000 S2 Reagent Kit (200 cycles). The average sequencing depth is 36.4 million reads per human macrophage sample.

### RNA-seq data pre-processing

FASTQ files generated by sequencing mouse paws were pre-processed by RNA-Seq Analysis Snakemake Workflow (RASflow) [61]. Briefly, raw read quality was evaluated using FastQC [62]. Reads were trimmed for adaptors and sequence quality using Trim Galore [63]. Trimmed reads were aligned to mouse GRCm39 reference genome using HISAT2 [64], and reads were counted by using featureCounts [65].

FASTQ files generated by sequencing human macrophages were imported into ArrayStudio (Qiagen). Briefly, raw data quality control was performed and then a filtering step was applied to remove reads corresponding to rRNAs as well as reads having low quality score or shorter than 25 nt. Reads were further mapped to the Human genome B38 using Omicsoft sequence aligner (OSA, version 4) [66] and quantified using Ensembl.R96 gene model. Paired reads were counted at the gene level.

For the public RNA-seq dataset on human synovial biopsies (GSE89408), raw counts at the gene level were extracted from HumanDisease_B38_GC33 of DiseaseLand (Qiagen). Briefly, DiseaseLand contains curated publicly available RNA-seq and expression array data, where all samples have been processed through the same pipeline to allow for cross project comparison. For HumanDisease_B38_GC33, alignment of FASTQ files was done using OSA to Human genome B38. The average sequencing depth is 34.1 million reads per human synovial sample.

### Statistical analysis on RNA-seq data

The statistical test on RNA-seq raw count data was performed by DESeq2 [67]. Depending on the respective study design and sequencing depth, genes that have >1 CPM (counts per million) in at least 20% of the mouse samples, at least 50% of the human macrophage samples, or at least 5% of the human synovium samples were included for analysis. For data on mouse samples, a time series differential analysis was done by the likelihood ratio test (LRT) in DESeq2, whereas for data on human macrophages and synovium (GSE89408), Wald test was used. The p-values attained by hypothesis testing in DESeq2 were corrected for multiple testing using the FDR method. Significant differentially expressed genes were defined by cutoffs of FDR < 0.05 and fold change > 1.5 in up or down direction. Genes that cannot be mapped to Entrez IDs, like novel pseudogenes, were removed from the list of differentially expressed genes. Furthermore, variance stabilizing transformation (VST) was applied to count data prior to PCA, and then the top 500 most variable genes were subjected to prcomp function (PCA) in the default stats package in R (v4.0.0).

### Gene set enrichment analysis

Gene set enrichment analysis was performed with the up-to-date biological process gene sets from the Gene Ontology database [26] using fgsea [68]. Genes were ranked in a decreasing order according to their fold changes, or to their loadings in a MOFA factor, and this ranking was used as the input to fgsea. By default, 1,000 permutations were done to calculate statistical significance. FDR-corrected p-value below 0.05 was considered as significant for a gene set.

### MOFA model training

MOFA model training was done on 171 mouse subjects that were profiled with both paw transcriptome and plasma metabolomics using MOFA2 package [31, 69]. Gene-level RNA-seq counts were pre-filtered and VST transformed using DESeq2. Metabolite intensities were log transformed. The inputs of MOFA model training comprised of the top 5,000 most variable genes and all 929 metabolites. Package default values were used for the training parameters, except for num_factors = 25, drop_factor_threshold = 0.02 and convergence_mode = slow. The latent factors and feature loadings were extracted from the best trained model with build-in functions of MOFA2.

### IPA upstream regulator analysis

Ingenuity Pathway Analysis (IPA, Qiagen) was used for prediction of upstream regulators from sets of identified differentially expressed genes [70]. The overlap p-values were calculated by the right-tailed Fisher’s exact test, and B-H adjusted for multiple testing.

### Other statistical analyses

For the associations between mouse groups (CIA, control) and phenotype features (scores, spleen weights, concentrations of cytokine or chemokine), two-tail Welch t tests was used. For the associations between mouse scores and factor values, one-way ANOVA was used. Both tests were done using base R package (v4.0.0).

### Visualization

The plots that depict study data and statistics were created with ggplot2 [71], pheatmap [72] or ggVennDiagram [73]. The schematic plots were created with BioRender.com.

## Data availability

The RNA sequencing data on CIA mouse paws and LPS-stimulated human macrophages have been deposited at Gene Expression Omnibus (GEO) database, under the accession numbers GSE193335 and GSE193336 respectively. The corresponding author will provide reviewer tokens for data access upon request, and the data will be publicly available as of the date of publication.

## Code availability

The code generated during the study are available from the corresponding author on reasonable request.

## Acknowledgements

We thank members of the Type 1/17 Immunology Cluster in Sanofi for technical support on the mouse experiments and Constanze Holz (Sanofi DMPK-Metabolism/tMSI) for technical support on the tissue MALDI-MS imaging. We are grateful to Annick Peleraux (Sanofi Biomarker Statistics) and Stephanie Pierson (Sanofi Pasteur/Digital R&D/Bioinformatics) for implementing the RASflow pipeline on the enterprise cloud computing environment. We thank Sandrine Tolou (Sanofi Translational Sciences) for the support on RNA sequencing of human macrophages and Mark Magid (Sanofi Precision Oncology) for depositing sequencing data into GEO database.

## Author contributions

N.B., C.A. and L.L. conceived and designed research studies. L.L., J.F., B.M., B.W., E.Z., G.T. and M.D. performed the experiments. L.L. and C.R. did the bioinformatic analyses. L.L. and N.B. wrote the manuscript, and all authors revised it. N.B. provided supervision and is the guarantor of this work.

