## Supplementary Document for "Multi-omics profiling of collagen-induced arthritis mouse model reveals early metabolic dysregulation via SIRT1 axis"

**Supplementary Table 1. Sample sizes for multi-omics experiments on CIA mice.**

| Timepoint | Treatment | Paw_RNAseq | Plasma_Metabolomics | Paw_MALDI-MS |
|---|---|---|---|---|
| 2 days | CIA | 18 | 18 | NA |
| 2 days | Ctrl | 6 | 6 | NA |
| 2 weeks | CIA | 15 | 17 | 6 |
| 2 weeks | Ctrl | 6 | 6 | 5 |
| 3 weeks | CIA | 14 | 18 | NA |
| 3 weeks | Ctrl | 6 | 6 | NA |
| 4 weeks | CIA | 16 | 17 | 6 |
| 4 weeks | Ctrl | 5 | 6 | 5 |
| 6 weeks | CIA | 17 | 18 | 6 |
| 6 weeks | Ctrl | 5 | 6 | 5 |
| 7 weeks | CIA | 17 | 17 | NA |
| 7 weeks | Ctrl | 5 | 5 | NA |
| 8 weeks | CIA | 16 | 16 | NA |
| 8 weeks | Ctrl | 5 | 5 | NA |
| 10 weeks | CIA | 15 | 16 | 6 |
| 10 weeks | Ctrl | 5 | 5 | 5 |

**Supplementary Table 2. Top 20 activated and top 20 inhibited upstream regulators from IPA on human synovium RNA-seq data.**

One exception here is that SIRT1 was not among the top 20 inhibited upstream regulators when considering its z score, but it was still significant at the stage of early RA.

| Upstream Regulators | Molecule Type | zScore. Arthralgia | BH_pval.Arthralgia | zScore. UA | BH_pval. UA | zScore. Early RA | BH_pval. Early RA |
|---|---|---|---|---|---|---|---|
| **lipopolysaccharide** | chemical drug | 4.95 | 0.014537887 | 8.998 | 4.11E-07 | 9.43 | 5.49E-14 |
| **IL1B** | cytokine | 4.907 | 0.021379927 | 7.602 | 2.52E-07 | 7.375 | 4.66E-10 |
| **TNF** | cytokine | 2.958 | 0.001881826 | 7.167 | 4.63E-08 | 7.269 | 7.53E-11 |
| **IFNG** | cytokine | 3.562 | 7.87E-05 | 5.294 | 1.98E-07 | 6.855 | 5.62E-13 |
| **poly rI:rC-RNA** | biologic drug | NA | 1 | 6.346 | 0.001285 | 6.85 | 1.47E-05 |
| **CSF2** | cytokine | 2.991 | 0.038963384 | 5.762 | 2.34E-05 | 6.22 | 4.30E-12 |
| **Interferon alpha** | group | 3.072 | 0.095024112 | 4.657 | 0.001586 | 6.22 | 1.21E-09 |
| **IL6** | cytokine | 3.391 | 0.015684944 | 6.085 | 6.48E-07 | 5.708 | 7.73E-16 |
| **OSM** | cytokine | 3.287 | 0.149468491 | 5.45 | 0.004161 | 4.29 | 0.005606 |
| **TLR9** | transmembrane receptor | NA | 1 | 4.996 | 0.008627 | 5.441 | 4.76E-07 |
| **PTPRR** | phosphatase | 5.379 | 7.87E-05 | 4.264 | 0.00075 | 3.162 | 0.194891 |
| **IL33** | cytokine | 4.014 | 0.01118701 | 4.799 | 3.81E-06 | 5.364 | 8.78E-10 |
| **MYD88** | other | NA | 1 | 5.35 | 0.001759 | 5.123 | 0.000104 |
| **CD28** | transmembrane receptor | 2.21 | 0.149194061 | 4.904 | 0.032463 | 5.301 | 3.41E-09 |
| **RELA** | transcription regulator | 3.682 | 0.028405697 | 5.019 | 1.58E-05 | 5.288 | 6.19E-09 |
| **NFkB (complex)** | complex | 2.115 | 0.000412795 | 4.429 | 1.40E-06 | 5.241 | 1.58E-12 |
| **IL2** | cytokine | 2.759 | 0.039192814 | 4.569 | 0.000191 | 5.225 | 4.14E-16 |
| **TNFSF12** | cytokine | 4.909 | 0.000165264 | 5.183 | 2.87E-06 | 4.81 | 2.50E-07 |
| **AHR** | ligand-dependent nuclear receptor | 3.531 | 0.055169976 | 3.275 | 0.000616 | 5.024 | 1.20E-09 |
| **TLR7** | transmembrane receptor | NA | 1 | 4.561 | 0.032646 | 5.018 | 3.37E-06 |
| **CITED2** | transcription regulator | NA | 1 | -5.331 | 0.018805 | -5.151 | 0.000432 |
| **Firre** | other | -4.914 | 1.24E-05 | -4.69 | 6.73E-05 | -3.742 | 0.006452 |
| **Tgf beta** | group | -4.659 | 0.000107735 | -1.974 | 2.42E-06 | -2.674 | 1.01E-09 |
| **PPARG** | ligand-dependent nuclear receptor | -2.987 | 0.464494386 | -4.648 | 0.037067 | -2.719 | 0.01186 |
| **ZFP36** | transcription regulator | -4.258 | 0.017980268 | -4.383 | 1.98E-07 | -4.605 | 6.57E-08 |
| **NUPR1** | transcription regulator | -1.921 | 0.394578913 | -2.06 | 0.235364 | -4.371 | 0.000475 |
| **TP53** | transcription regulator | -4.26 | 0.304135769 | -3.912 | 0.067025 | -3.736 | 0.0039 |
| **EHMT1** | transcription regulator | -4.082 | 0.007006118 | -3.578 | 0.005132 | -1.678 | 0.009123 |
| **Irgm1** | other | NA | 1 | -3.063 | 0.030323 | -4.022 | 8.88E-05 |
| **IL1RN** | cytokine | -3.201 | 0.026427214 | -3.626 | 0.000645 | -4.018 | 8.91E-05 |

| | | | | | | | |
|---|---|---|---|---|---|---|---|
| **SP1** | transcription regulator | -3.953 | 0.017617377 | -1.95 | 0.00036 | 0.311 | 0.002874 |
| **SS18** | transcription regulator | -3.742 | 0.001326942 | -2.111 | 0.003711 | -1.508 | 0.000292 |
| **CTLA4** | transmembrane receptor | -2.797 | 0.038563448 | -3.089 | 0.001759 | -3.659 | 4.74E-09 |
| **DAP3** | other | -3.317 | 2.52E-06 | -3.317 | 6.55E-08 | -3.606 | 1.58E-13 |
| **LONP1** | peptidase | -2.887 | 0.334364038 | -1.678 | 0.027521 | -3.606 | 0.009718 |
| **ESRRA** | transcription regulator | -3.604 | 0.031263041 | NA | 1 | -0.077 | 0.13618 |
| **CDKN1A** | kinase | NA | 1 | -3.054 | 0.030854 | -3.509 | 5.81E-06 |
| **LILRB4** | other | -3.211 | 0.022828467 | -3.211 | 0.00304 | -3.497 | 8.08E-06 |
| **NORAD** | other | -3.45 | 0.021379927 | -1.706 | 0.045173 | -2.121 | 0.02357 |
| **DUSP1** | phosphatase | -3.151 | 0.040947334 | -3.435 | 2.87E-06 | -3.441 | 6.00E-06 |
| **SIRT1** | transcription regulator | NA | 1 | NA | 1 | -1.36 | 0.043221 |

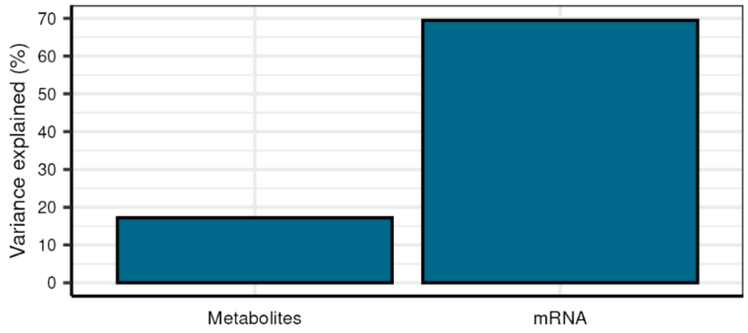


**Fig. S1: Cumulative variance explained by all factors in each data modality.**

MOFA identified in total 6 factors, based on the criterion that a factor should account for > 2% of the variance in at least one omics dataset. All the factors together explained approximately 20% of variance in metabolomics dataset, and approximately 70% of variance in transcriptomics dataset.
